# Role of the SAF-A/HNRNPU ATPase and RGG domains in X chromosome inactivation, nuclear dynamics, transcription, splicing, and cell proliferation

**DOI:** 10.64898/2025.12.15.694381

**Authors:** Judith A. Sharp, Rachael Thomas, Emily Sparago, Wei Wang, Michael D. Blower

**Affiliations:** Department of Biochemistry and Cell Biology, Chobanian and Avedisian School of Medicine, Boston University 72 E. Concord St, K112, Boston, MA 02118 USA

## Abstract

The SAF-A/HNRNPU gene encodes an abundant nuclear protein conserved throughout vertebrates, and is mutated in individuals with HNRNPU syndrome, a neurological human disease. SAF-A is important for maintaining lncRNA localization, splicing, and gene expression state. The mechanistic role of SAF-A in each of these processes is coordinated by one or more of its functional domains, which include an N-terminal SAP domain, a central ATPase domain, and a series of C-terminal RGG repeats embedded in a low-complexity region. The SAP domain and RGG repeats define two nucleic acid interaction domains, with both capable of binding DNA or RNA. Here we use an allelic reconstitution strategy to investigate the role of the SAF-A ATPase domain and RGG repeats. We show that both the ATPase and RGG repeats control SAF-A nuclear dynamics, and present genetic evidence that SAF-A interacts with nascent transcripts through the RGG repeats. The SAF-A ATPase domain and RGG repeats were also required for maintaining XIST RNA and facultative heterochromatin marks on the inactive X chromosome, with distinct effects of mutations that block ATP binding and ATP hydrolysis. Analysis of transcriptome datasets revealed that the SAF-A ATPase domain and RGG repeats are both required for proper mRNA splicing, but not for gene expression. Importantly, we found that like the SAP domain, the SAF-A ATPase domain and RGG repeats are required for cell proliferation. Collectively, our findings highlight the importance of the SAF-A SAP, ATPase and RGG domains in essential functions of nuclear biology. These analyses will therefore inform our understanding of the disease state in HNRNPU syndrome.

## Introduction

RNA is important for forming compartments in the nucleus to regulate essential functions, such as RNA processing and gene expression [1–5]. RNA and RNA-binding proteins localize near active chromatin to organize a network of transcription factors and other chromatin regulators. The proper assembly of transcription-associated nuclear structures is important for essential nuclear functions and to prevent neurological disease [6, 7].

The SAF-A/HNRNPU protein is a highly abundant nuclear RNA-binding protein that is essential in mice [8, 9] and cancer cells [10]. *De novo* mutations in SAF-A in humans or mice lead to a variety of neurological defects [11–13]. SAF-A was originally described as a factor that binds to hnRNA and attaches RNA to nuclear scaffold attachment regions [14–16]. SAF-A has been implicated in regulation of mRNA splicing [8, 11, 12, 17], gene expression [11, 12], nuclear structure [18, 19], and tethering of the XIST and FIRRE lncRNAs to chromatin [20–23]. The SAF-A protein has several conserved functional domains, including an N-terminal SAP domain, a D/E-rich acidic region, a SPRY domain of unknown function, a central ATPase domain, and a C-terminal RGG domain involved in binding RNA and DNA (Figure 1A; [24]). We recently demonstrated that the SAP domain and SAP domain phosphorylation is important for cell viability, XIST RNA localization, and controlling protein dynamics [25]. At present, very little is known about the roles of the ATPase and RGG domains within cells.

**Figure 1.**
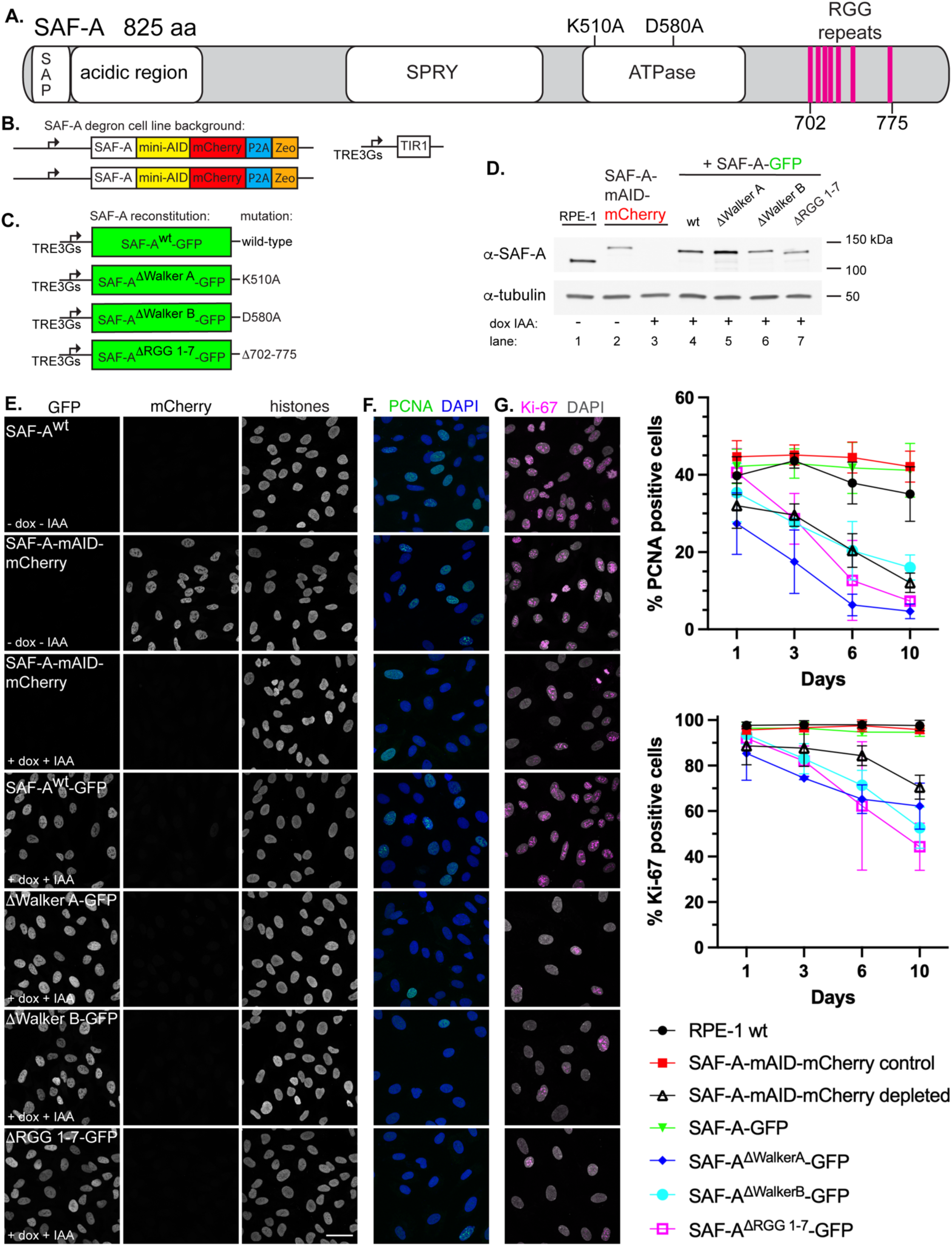
The ATPase and RGG domains are required for cell proliferation. A. Schematic depicting full-length SAF-A (isoform a, 825 amino acids) with domains drawn to scale. The position of the seven RGG repeats is marked between residues 702-775. B. Design of the SAF-A-AID-mCherry degron to replace the endogenous SAF-A genes in RPE-1 cells. C. SAF-A-GFP tagged transgenes used to reconstitute either wild-type SAF-A, or versions of SAF-A bearing point mutations in the Walker A or Walker B elements in the ATPase domain, or a deletion of residues 702-775 containing RGG repeats 1-7. D. Western blot analysis of SAF-A cell lines to monitor expression of SAF-A-AID-mCherry (α-RFP panel) and SAF-A-GFP (α-GFP panel) in cell lines. Tubulin was used as a loading control. Lane 1: untagged RPE-1 parental cell line. Lane 2: SAF-A degron cell line, either without (lane 2), or with (lane 3) doxycycline and IAA. Lanes 4-7: cells expressing SAF-A-GFP transgenes, treated with doxycycline and IAA. Molecular weight markers are indicated to the right of the panel. E. Immunofluorescence of cells expressing tagged SAF-A, as indicated by the panel insets. Treatment with doxycycline and IAA as indicated led to complete depletion of SAF-A-AID-mCherry in the vast majority of cells. Integrated SAF-A-GFP transgenes showed uniform expression in the cell population after induction. Bar, 40 μm. F-G. PCNA and Ki-67 immunofluorescence to monitor cell proliferation. Cells were treated with doxycycline and IAA to induce endogenous SAF-A depletion and SAF-A-GFP transgene expression, and were compared to control cell populations. Cells were treated for 1, 3, 6, or 10 days as indicated. SAF-A expression is the same as the panel insets shown in E. All images in panels E-G are rendered as a maximum projection of a 3D stack. H. Quantitation of cell populations with PCNA- and Ki-67-positive cells expressed as percent of the total population. 300 cells were scored for PCNA and Ki-67 immunofluorescence in two biological replicates; the average and standard deviation is shown.

Among RNA binding motifs, the RGG domain is the second most common in the human genome [26]. RGG domains consist of a series of RGG or RG repeats interspersed with other amino acids. RGG domains can be used to interact with RNAs and also for protein-protein interactions [27]. The C-terminus of SAF-A contains seven RGG elements embedded within a larger low-complexity domain, featuring a Q/N-rich region [21, 28]. *In vitro* binding studies point to the first six RGG repeats as being neither necessary nor sufficient for RNA binding [29, 30]. Interestingly however, deletion of RGG repeats 1-7 causes dramatically reduced RNA binding *in vitro* [29]. Overall, the RNA binding activity contained within the SAF-A C-terminus interacts broadly with RNA and shows very little sequence specificity [26, 30].

SAF-A is localized to the inactive X chromosome (Xi) in female cells and interacts directly with the XIST lncRNA required for assembling facultative heterochromatin on the Xi [31]. SAF-A is required for the proper localization of XIST RNA to the Xi territory [21]. Previous analyses of SAF-A mutations found that deleting the entire C-terminus of SAF-A results in defective XIST RNA localization, as does a smaller deletion of low-complexity sequences downstream of the seven RGG elements [21, 28]. One study noted that an internal deletion of the first six RGG elements had no effect on XIST RNA localization [32]. However, it is not presently known whether the RNA binding activity defined by SAF-A RGG repeats 1-7 affects XIST RNA localization, or other SAF-A nuclear functions.

In addition to a RGG domain, SAF-A contains a poorly characterized, conserved central ATPase domain. Initial structural homology modeling of the SAF-A ATPase domain suggested a strong similarity to the AAA+ domain of RecA, and it was proposed that SAF-A uses a combination of RNA binding and ATP-binding to form a helical polymer [19]. However, a follow-up study showed that the SAF-A ATPase domain is more closely related to a P-loop NTPase domain and is unlikely to use this domain to form higher order oligomers [33]. *In vivo*, mutation of the SAF-A Walker-A or Walker-B motifs resulted in changes in SAF-A complex formation and the ability of SAF-A to decondense chromatin, suggesting that the ATPase domain is important for SAF-A function [19]. However, the molecular function of the SAF-A ATPase domain is unknown and it is not clear if ATP binding and hydrolysis are important for the many functions that have been attributed to SAF-A.

In this study we present phenotypic analysis of cell lines either lacking SAF-A expression, or expressing mutations that prevent ATP binding, prevent ATP hydrolysis, or dramatically reduce RNA binding. We found that the ATPase domain and RGG repeats 1-7 were important for SAF-A function in XIST RNA localization and activity, SAF-A dynamic nuclear mobility, interaction with nascent RNA, normal mRNA splicing, and cellular proliferation. In contrast, the ATPase and RGG domains were not required for normal gene expression. Taken together, our data suggest that the ATPase and RGG domains work together in a dynamic cycle to promote the majority of SAF-A functions in the interphase nucleus.

## Results

### The SAF-A ATPase and RGG domains are required for cell proliferation

To investigate the role of the ATPase and RGG domains in SAF-A nuclear functions, we utilized a depletion-reconstitution strategy that we had used previously to study the SAP domain in diploid, karyotypically stable human RPE-1 cells ([25, 29]; Figure 1B-C). In this system, both endogenous SAF-A genes are tagged at the 3’ end with an auxin-inducible degron sequence and the mCherry fluorescent protein and are expressed from the native promoter (SAF-A-AID-mCherry; Figure 1B). The TIR1 E3 ligase that mediates auxin-dependent protein degradation is also stably integrated and expressed from a doxycycline-inducible promoter. Addition of doxycycline and 3-indole acetic acid (IAA) causes the simultaneous induction of the TIR1 E3 ligase and degradation of SAF-A-AID-mCherry [25, 29, 34]. To quantitate the extent of SAF-A depletion after 24 hours of drug treatment, we analyzed SAF-A-AID-mCherry expression in cell populations and cell extracts (Figure 1D-E). Under these conditions, we observed 98% of cells showed complete depletion of SAF-A-AID-mCherry after addition of doxycycline and IAA (Figure 1E and Supplemental Figure 1A). Western blot analysis confirmed depletion of the degron allele (Figure 1E, lanes 2-3 and Supplemental Figure 1B).

We then used lentiviral integration to introduce doxycycline-inducible, GFP-tagged transgenes to analyze SAF-A function with the following constructs (Figure 1C): 1) full-length, wild-type SAF-A (SAF-A^wt^-GFP), 2) a mutation in the SAF-A ATPase domain that blocks ATP binding (K510A; SAF-A^ΔWalker^ ^A^-GFP), 3) a mutation in the Walker B motif predicted to block ATP hydrolysis (D580A; SAF-A^ΔWalker^ ^B^-GFP), and 4) a deletion encompassing the seven RGG repeats in the RNA binding domain (Δ702-775; SAF-A^ΔRGG^ ^1-7^-GFP). Immunofluorescent detection of GFP and mCherry in these cell lines confirmed the simultaneous depletion of endogenous SAF-A-mAID-mCherry and inducible expression of SAF-A-GFP alleles throughout cell populations, after 24 hours of drug treatment (Figure 1E). Western blot analysis confirmed that the reconstituted cell lines expressed the SAF-A-GFP alleles as the only source of SAF-A expression, and that all SAF-A-GFP alleles were expressed at levels similar to the endogenous protein (Figure 1D, lanes 4-7, Supplemental Figure 1B). Our studies have confirmed that SAF-A^wt^-GFP behaves comparably to the native, untagged protein in all functional assays tested thus far, including cell viability, gene expression, epigenetic features of the inactive X chromosome, and splicing [25, 29, 34]. These cell lines therefore represent a defined system to analyze the specific contributions of the ATPase and RGG domains to SAF-A nuclear functions.

SAF-A is an essential gene required for embryonic viability and cell proliferation [8, 9]. Our previous analysis identified that the N-terminal SAP domain and SAP domain phosphorylation has a critical role in supporting the proliferative capacity of cell populations [25]. To test whether the ATPase or RGG domains have a role in maintaining cell division, we analyzed the number of cycling cells using PCNA and Ki-67 as markers for cell proliferation. Over the 10 day time course of the experiment, SAF-A depleted cells showed reduced numbers of PCNA- and Ki-67-positive cells, indicating that cells were exiting the cell cycle in response to the loss of SAF-A. Loss of cell proliferation was completely rescued by expression of SAF-A^wt^-GFP. In contrast, cells expressing SAF-A^ΔWalker^ ^A^-GFP, SAF-A^ΔWalker^ ^B^-GFP, and SAF-A^ΔRGG^-GFP also showed a reduced percentage of PCNA- and Ki-67-positive cells, and exited the cell cycle with kinetics similar to the null condition. From these data we conclude that in addition to the SAP domain, the ATPase and RGG domains are each independently required to maintain normal levels of cell proliferation, and are therefore important for essential functions of SAF-A.

### SAF-A exhibits rapid nuclear dynamics and interacts with nascent transcripts using the RGG domain

To test whether the ATPase or RGG domains play a role in SAF-A nuclear dynamics in living cells, we performed FRAP analysis in cells expressing wild-type and mutant SAF-A-GFP transgenes (Figure 2). SAF-A^wt^-GFP exhibited rapid nuclear dynamics with a t_1/2_ of recovery of 2.58 seconds (Figure 2A-C). An average of only 18% of SAF-A^wt-^GFP molecules were immobile during the 30 second experiment, thus representing a high degree of nuclear mobility [25]. Next, we compared SAF-A mobility in cells expressing mutations of the ATPase and RGG domains: SAF-A^ΔWalkerA^-GFP, SAF-A^ΔWalkerB^-GFP, and SAF-A^ΔRGG^ ^1-7^-GFP. Both ATPase domain mutations decreased the t_1/2_ of recovery to approximately 2.0 seconds with a corresponding decrease in immobile fraction, representing more rapid nuclear motion than the wild-type protein (Figure 2B-C). These data suggest that ATP binding and hydrolysis make a modest contribution to overall SAF-A nuclear dynamics. In contrast, deletion of the RGG domain resulted in a dramatic increase in mobility, with the t_1/2_ of recovery decreasing to ∼1.2 seconds and causing a complete loss of the immobile fraction. This result demonstrates that the RGG domain is required for the majority of SAF-A interactions in the nucleus, while the ATPase domain plays a more minor role.

**Figure 2.**
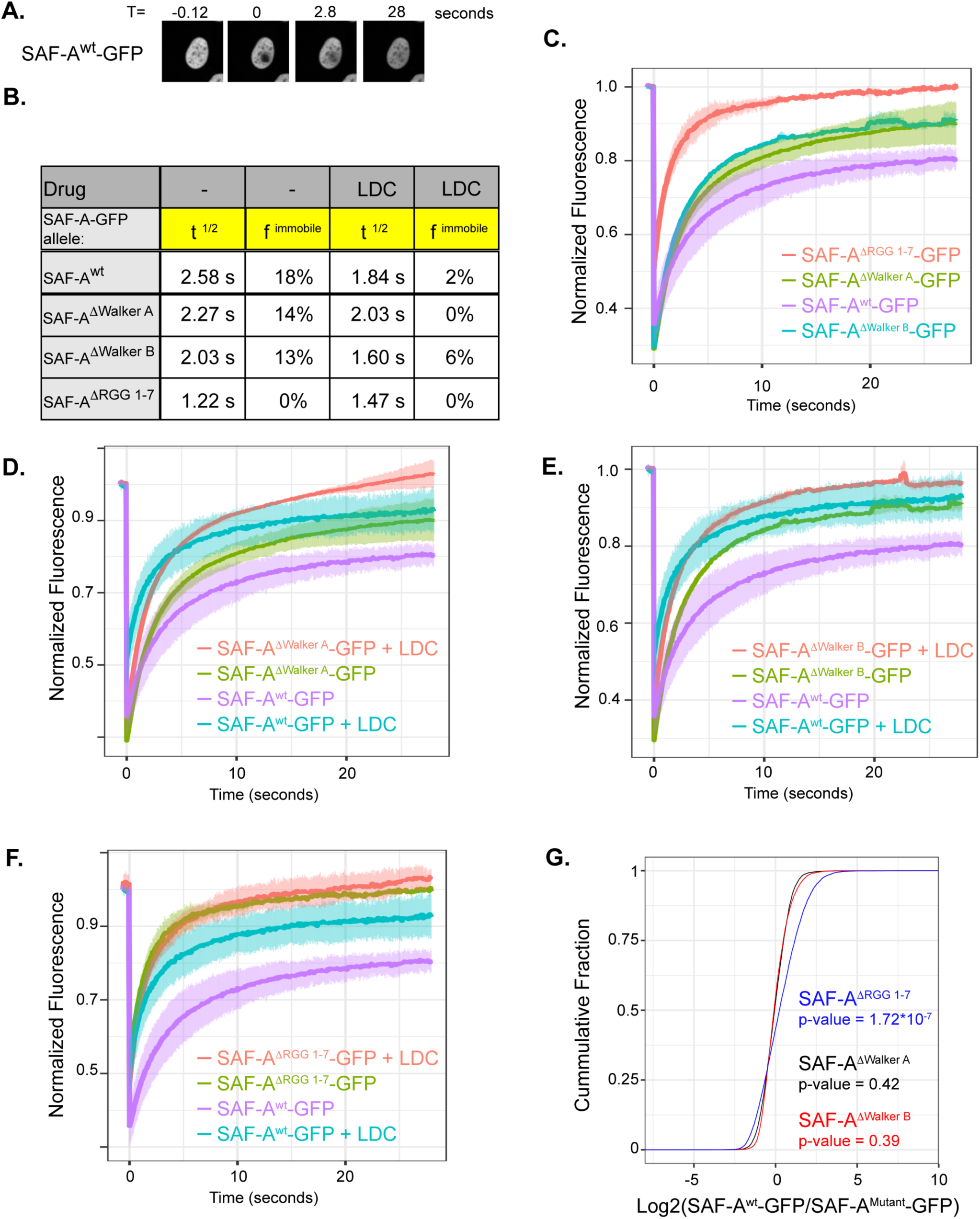
The SAF-A RGG and ATPase domains are important for nuclear dynamics. A. Images from a typical FRAP experiment using SAF-A^wt^-GFP. B. Table of the t^1/2^ recovery time and the immobile fraction for all SAF-A-GFP proteins, with and without transcriptional inhibition. C-F. FRAP recovery curves for SAF-A^wt^-GFP, SAF-A^ΔWalker-A^-GFP, SAF-A^ΔWalkerB^-GFP, and SAF-A^ΔRGG^-GFP, with and without transcriptional inhibition (LDC). The standard deviation of recovery time is indicated in light colored error bars. G. SAF-A^wt^-GFP, SAF-A^ΔWalker-A^-GFP, SAF-A^ΔWalkerB^GFP, and SAF-A^ΔRGG^-GFP were immunoprecipitated and co-precipitated RNAs were sequenced and normalized to *Drosophila* spike-in control RNA. Enrichment of RNA in all mutants was compared to WT and is plotted as a Cumulative Distribution Function.

Recent work demonstrated that SAF-A localization and activity is responsive to nuclear transcription [1, 19], and our recent work demonstrated that transcriptional inhibition leads to a dramatic increase in SAF-A mobility [25]. To determine whether the ATPase and RGG domains contribute to SAF-A nuclear dynamics in the absence of transcription, we performed FRAP after inhibition of transcriptional elongation using the CDK9 inhibitor LDC000067 (LDC) [35]. Control experiments confirmed that acute inhibition of CDK9 resulted in a strong decrease in nascent transcription [25].

Interestingly, SAF-A^wt^-GFP and both ATPase domain mutants exhibited increased nuclear mobility in the presence of LDC (Figure 2B, 2D-E), suggesting that the ATPase domain is not important for the interaction with nascent transcripts. In contrast, SAF-A^ΔRGG^ ^1-7^-GFP mobility was largely unaffected by transcriptional inhibition. These data demonstrate that the main binding site for SAF-A in the nucleus is nascent RNA, and that SAF-A binds nascent RNA using the RGG domain (Figure 2F). Taken together, these data demonstrate that SAF-A is a highly dynamic protein that interacts with nascent transcripts through the RGG domain.

Previously, we showed that a deletion of the seven RGG repeats resulted in a ∼10-fold reduction of RNA binding *in vitro* [29]. To determine whether SAF-A^ΔRGG^ ^1-7^-GFP or mutations of the ATPase domain caused a reduction in the global interaction of SAF-A with RNA *in vivo*, we immunoprecipitated all SAF-A-GFP species and sequenced co-purifying RNA . In brief, SAF-A-GFP transgenes were expressed in cells depleted for endogenous SAF-A-AID-mCherry as described above. We prepared cell extracts and used a GFP antibody to immunopurify wild-type and mutant versions of SAF-A-GFP. We then spiked in total *Drosophila* RNA as a normalization control, in order to compare RNA binding between genotypes after sequencing (Materials and methods). We found that mutation of the ATPase domain had no effect on RNA association with SAF-A (Figure 2G). In contrast, deletion of the RGG repeats 1-7 resulted in a dramatic reduction in global RNA binding, consistent with our *in vitro* RNA binding results (Figure 2G)[29]. We conclude that ATP binding and hydrolysis are not required for SAF-A interaction with RNA, and that the majority of SAF-A RNA binding occurs through the RGG domain.

### SAF-A targeting to the Xi is independent of the ATPase and RGG domains

SAF-A localizes to the Xi chromosome territory as defined by the Barr body and is modestly enriched on the Xi in interphase cells relative to other nuclear regions [25]. To determine which domains of SAF-A are required for Xi localization, we performed live cell imaging of cells expressing SAF-A^wt^-GFP, SAF-A^ΔSAP^-GFP, SAF-A^ΔWalker^ ^A^-GFP, SAF-A^ΔWalker^ ^B^-GFP, and SAF-A^ΔRGG^ ^1-7^-GFP (Figure 3). As expected, SAF-A^wt^-GFP showed Xi enrichment, while SAF-A^ΔSAP^-GFP was completely excluded from the Xi territory, confirming our previous finding that the SAP domain is critical for Xi targeting of SAF-A (Figure 3A-B) [25, 36]. Both SAF-A^ΔWalker^ ^A^-GFP and SAF-A^ΔWalker^ ^B^-GFP were observed on the Xi, suggesting that neither ATP binding nor ATP hydrolysis are a prerequisite for SAF-A enrichment on the Xi (Figure 3C-D). Surprisingly, SAF-A^ΔRGG^ ^1-7^-GFP also showed Xi localization, indicating that the recruitment of SAF-A to the Xi is independent of the RNA binding activity contained within SAF-A residues 702-775 (Figure 3E-F). Therefore, for all SAF-A domains analyzed thus far, only the SAP domain is uniquely required for Xi targeting of SAF-A (this study, [25]).

**Figure 3.**
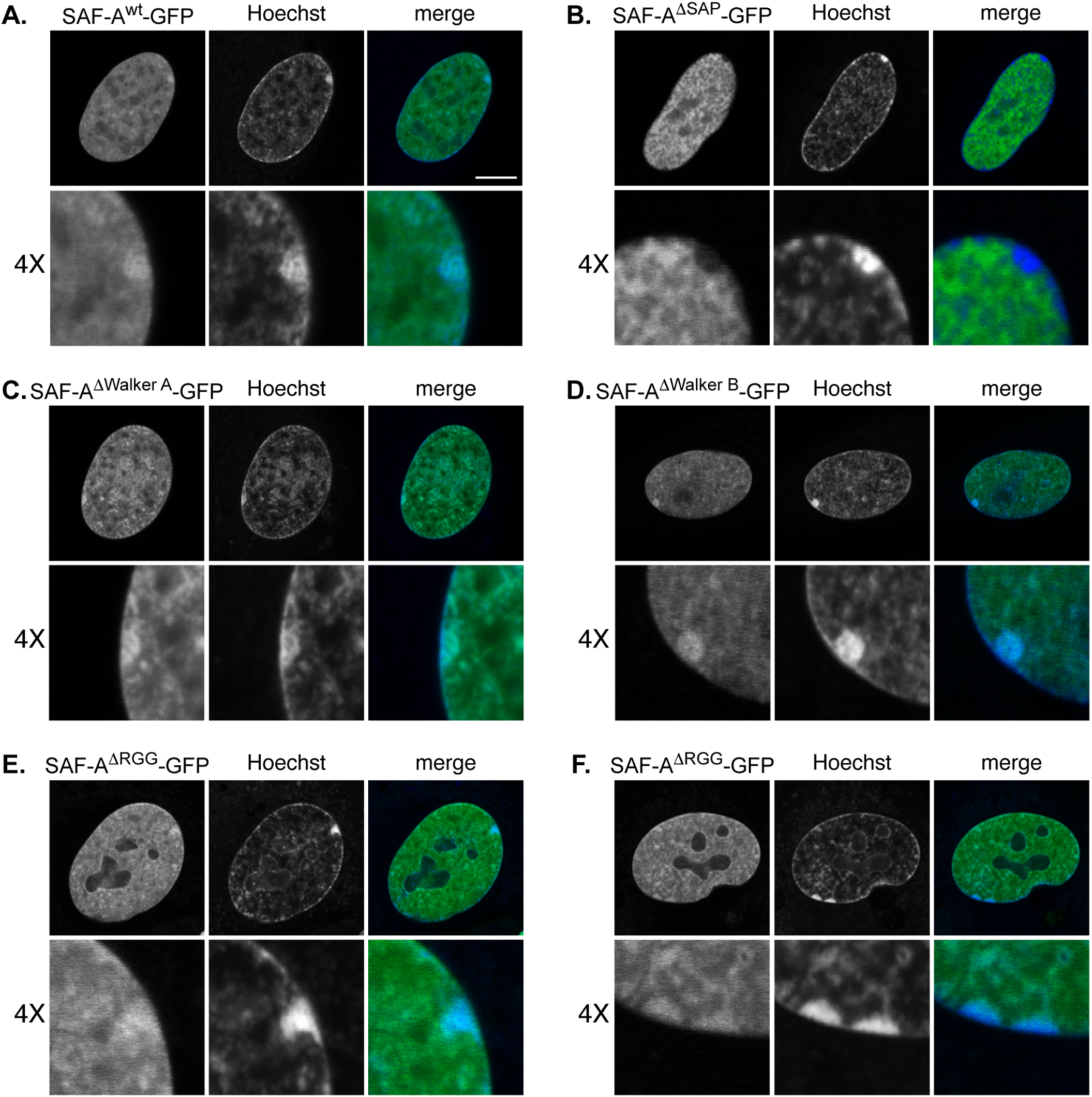
Recruitment of SAF-A to the Xi is independent of the ATPase and RGG domains. A-F. Live cell analysis of SAF-A^wt^-GFP and SAF-A domain mutations as indicated. Two examples of SAF-A^ΔRGG^ ^1-7^-GFP are shown. Cells were analyzed 24 hours after doxycycline and IAA treatment. Each image represents a single 0.2 mm slice. Bar, 10 μm. The proportion of cells showing the depicted localization patterns for each allele are as follows: SAF-A^wt^-GFP, 97% of cells; SAF-A^ΔSAP^ -GFP, 100% of cells; SAF-A^ΔWalker^ ^A^-GFP 100% of cells; SAF-A^ΔWalker^ ^B^-GFP 99% of cells; SAF-A^ΔRGG^ ^1-7^-GFP 100% of cells (n ≥ 50). At least two independent live imaging experiments for each genotype were performed.

A previous study reported that the modest enrichment of SAF-A in the Xi territory was disrupted by a deletion of SAF-A RGG repeats 1-6 in HEK293 cells [37], while we observe that SAF-A^ΔRGG^ ^1-7^-GFP is present on the Xi. We note that there are experimental differences between our assay and the methods used previously. In that study, the authors used immunofluorescence to detect SAF-A in fixed cell populations, while we are using live cell imaging. We have reported observing differences in SAF-A localization in fixed cell populations, suggesting there are rearrangements of SAF-A localization during formaldehyde crosslinking that do not occur in intact cells [25]. We have also observed that internal SAF-A mutations reduce the avidity of antisera [25]. It is possible that either of these experimental differences could explain the discrepancy in our findings.

### The SAF-A ATPase and RGG domains support XIST RNA localization to the Xi

SAF-A is required to maintain normal XIST RNA localization to the Xi in both human and murine female cells [21–23, 25, 28, 38], in a manner independent of XIST RNA gene expression [[25], Supplemental Figure 2A]. Previously, we reported that SAF-A depletion in our RPE-1 degron system leads to XIST RNA delocalization within one cell cycle, suggesting SAF-A is continuously required to maintain XIST RNA concentration in the Xi territory. Further, in order to accurately compare the impact of different SAF-A mutations on RNA localization, we reported a quantitative imaging method to count the number of XIST RNA foci per cell [25]. In the confocal microscope, cells with normal XIST RNA localization have 1-5 resolvable XIST RNA foci per cell, due to the close positioning of XIST RNA molecules within the Xi territory. SAF-A mutations causing a localization defect have a statistically significant increase in foci, due to the presence of XIST RNA molecules located distal to the Xi.

To test whether the SAF-A ATPase and RGG domains are required for XIST RNA localization, we treated cell lines for 24 hours to deplete endogenous SAF-A-AID-mCherry and induce SAF-A-GFP transgenes. We then performed XIST RNA FISH hybridization on fixed cell populations and acquired 3D optical stacks for ≥ 100 interphase nuclei per genotype in two independent experiments. Each image set was analyzed in batch to count XIST RNA foci using Fiji software [25]. Figure 4A depicts representative XIST RNA FISH patterns for each genotype; the quantitative analysis of XIST RNA foci per cell is shown as a violin superplot in Figure 4B [39]. Statistical comparisons between genotypes are listed in the figure legend.

**Figure 4.**
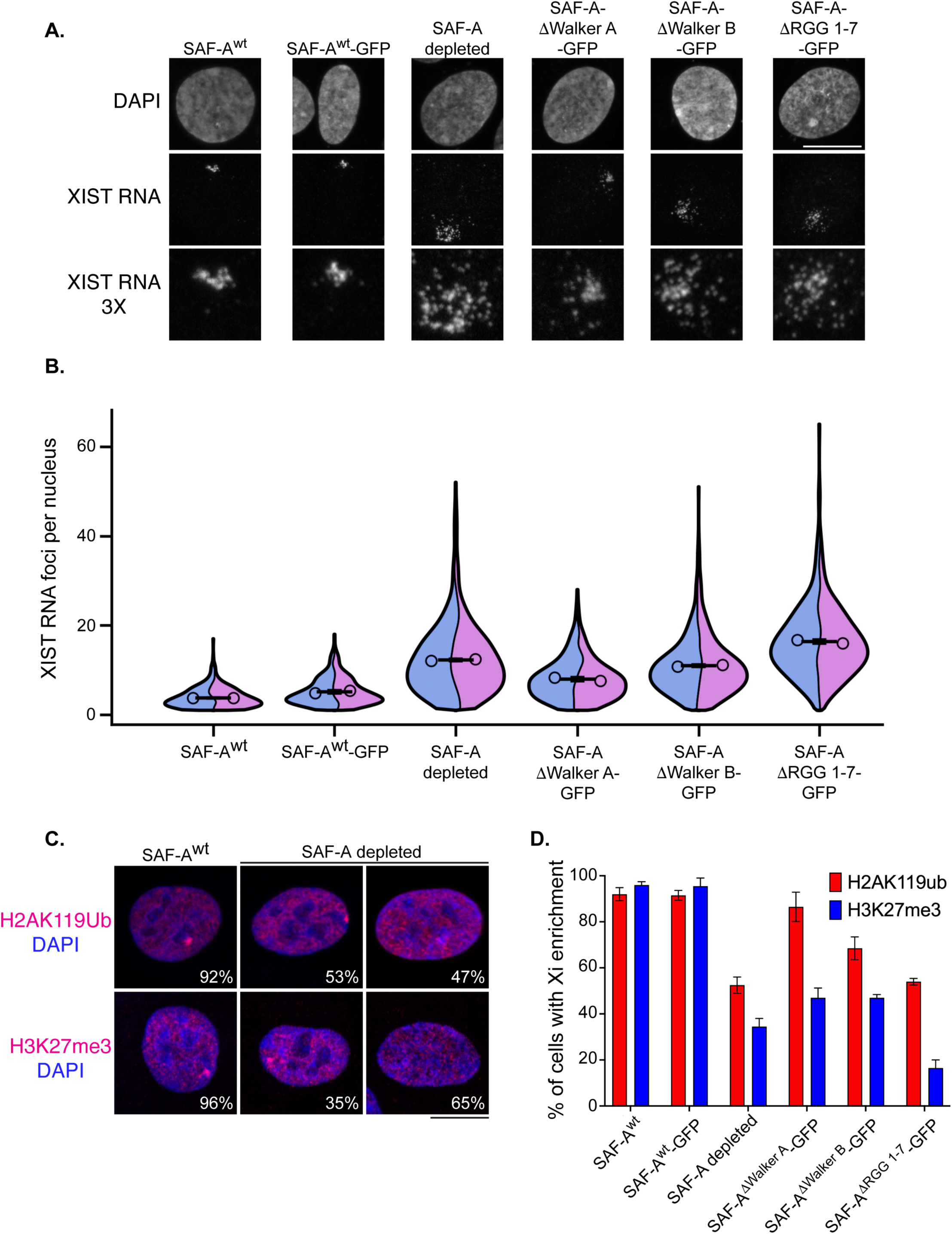
The ATPase domain and RGG 1-7 repeats impact XIST RNA localization and PRC1/PRC2-dependent histone modifications on the Xi. A. XIST RNA FISH and DAPI staining in RPE-1 cells, and cells expressing SAF-A-GFP transgenes, 24 hours after treatment with doxycycline and IAA. Images are rendered as a maximum projection of a 3D stack. Bar, 10 μm. B. Quantitative measurement of XIST RNA foci per cell. The term foci refers to the number of resolvable objects rather than individual molecules. Image stacks were acquired for at least 100 nuclei per genotype for two biological replicates and analyzed in Fiji software to count XIST RNA foci. Measurements are depicted as violin superplots. The average of each replicate is depicted as an open circle, whereas the average of both replicates is depicted as a horizontal line. The standard deviation of the two averages is shown. Cell n for quantitation of XIST RNA particles is: RPE-1 n = 193 and n = 211, SAF-A^wt^-GFP n = 137 and n = 236, SAF-A depleted 1 n = 122 and n = 220, SAF-A^ΔWalker-A^-GFP n = 135 and n = 193, SAF-A^ΔWalker^ ^B^-GFP n = 177 and n = 183, SAF-A^ΔRGG^ ^1-7^-GFP n = 159 and n = 180. Statistical comparison of number of XIST RNA particles was performed using a one-way ANOVA followed by Tukey’s tests with a Bonferroni correction. The *p*-value comparisons between SAF-A^wt^ and all other genotypes are as follows: SAF-A^wt^ versus SAF-A^wt^-GFP, ns. SAF-A^wt^ versus SAF-A depleted, *p* = 0.001. SAF-A^wt^ versus SAF-A^ΔWalker-A^-GFP, 0.088. SAF-A^wt^ versus SAF-A^ΔWalker^ ^B^-GFP, *p* = 0.003. SAF-A^wt^ versus SAF-A^ΔRGG^ ^1-7^-GFP, *p* = 0.001. C. Immunofluorescence of histone modifications H2AK119ub and H3K27me3 in RPE-1 cells and SAF-A depleted cells, 24 hours after treatment with doxycycline and IAA. Nuclei were stained with DAPI. Merged images are rendered as a maximum projection of a 3D stack. Bar, 10 μm. D. Quantitation of cell populations with H2AK119ub and H3K27me3 enrichment on the Xi. 100 cells were scored in two biological replicates; the average and standard deviation is shown. Statistical comparison was performed using a t-test to determine *p*-values for H2AK119ub enrichment. SAF-A^wt^ versus SAF-A^wt^-GFP, *p* = 0.860, ns. SAF-A^wt^ versus SAF-A^ΔWalker-A^-GFP, *p* = 0.380, ns. All other comparisons were statistically significant: SAF-A^wt^ versus SAF-A depleted, *p* = 0.007. SAF-A^wt^ versus SAF-A^ΔWalker^ ^B^-GFP, *p* = 0.028. SAF-A^wt^ versus SAF-A ^ΔRGG^ ^1-7^-GFP, *p* = 0.003. The same analysis was performed to determine *p*-values for H3K27me3 enrichment. SAF-A^wt^ versus SAF-A^wt^-GFP, *p* = 0.870, ns. All other comparisons were statistically significant: SAF-A^wt^ versus SAF-A depleted, *p* = 0.002. SAF-A^wt^ versus SAF-A^ΔWalker-A^-GFP, *p* = 0.004. SAF-A^wt^ versus SAF-A^ΔWalker^ ^B^-GFP, *p* = 0.001. SAF-A^wt^ versus SAF-A^ΔRGG^ ^1-7^-GFP, *p* = All images shown are projections of an optical stack of 0.2 μm slices.

Both native RPE-1 cells (SAF-A^wt^) and SAF-A^wt^-GFP control cells showed comparable numbers of XIST foci per nucleus, whereas the SAF-A depleted cells resulted in a 2-fold increase of XIST foci per nucleus (Figure 4B). Also of note is the presence of the long violin distribution in the SAF-A depleted samples, indicative of a significant percentage of cells with 10-50 XIST RNA foci per nucleus. Cells expressing SAF-A^ΔWalker^ ^A^-GFP had a relatively modest phenotype, suggesting ATP binding causes only mild perturbation of XIST RNA localization. In contrast, cells expressing either SAF-A^ΔWalker^ ^B^-GFP or SAF-A^ΔRGG^ ^1-7^-GFP showed a distribution of mislocalized XIST RNA foci similar to the null condition, arguing that ATP hydrolysis by SAF-A is important for XIST RNA localization, as is the RNA binding activity contained within the RGG repeats.

### The SAF-A ATPase and RGG domains are required for maintenance of Polycomb-dependent histone modifications on the Xi

The XIST RNP recruits the histone modification enzymes PRC1 and PRC2 to the Xi, resulting in the enrichment of H2AK119ub and H3K27me3 on Xi chromatin [40]. This process depends in part on SAF-A, since depletion of SAF-A results in the loss of these histone modifications on the Xi in a significant fraction of cells [21, 29]. Specifically, while wild-type RPE-1 cell populations have 92% of cells showing H2AK119ub Xi enrichment, we observed that in SAF-A depleted cell populations, only 53% of cells showed this pattern (Figure 4C-D). Similarly, RPE^wt^ cells had 96% of cells with H3K27me3 Xi enrichment, in contrast to only 35% of SAF-A depleted cells (Figure 4C-D). Cells expressing SAF-A^wt^-GFP had H2AK119ub and H3K27me3 Xi enrichment frequencies comparable to RPE-1 control cells. Cells expressing SAF-A^ΔWalker^ ^A^-GFP showed an unexpected pattern: 87% of cells had normal H2AK119ub Xi enrichment, but showed reduced H3K27me3 Xi enrichment, occurring in only 47% of cells. These data suggest that PRC1 recruitment and activity on the Xi is independent of SAF-A ATP binding. In contrast, normal PRC2 activity on the Xi requires SAF-A ATP binding. To our knowledge, this is the first example illustrating that PRC2 function on the Xi can be defective even when H2AK119ub Xi enrichment is intact. Cells expressing SAF-A^ΔWalker^ ^B^-GFP and SAF-A^ΔRGG^ ^1-7^-GFP alleles had reduced levels of both H2AK119ub and H3K27me3 modifications on the Xi, with SAF-A^ΔRGG^ ^1-7^-GFP cells having the lowest incidence of H3K27me3 (17% of cells) of all genotypes scored. These data show that ATP hydrolysis and RGG-dependent RNA binding by SAF-A both contribute to H2AK119ub and H3K27me3 Xi enrichment. Overall, the role of XIST RNA in maintaining Polycomb-dependent histone modifications on the Xi depends on both the SAF-A ATPase domain and RGG 1-7 repeats.

### SAF-A mutants do not affect global gene expression or gene expression on the Xi

To test whether mutation of the ATPase or RGG domains affects steady-state RNA levels in RPE-1 cells, we performed RNA-seq in cells expressing SAF-A^wt^-GFP, SAF-A^ΔWalker^ ^A^-GFP, SAF-A^ΔWalker^ ^B^-GFP, and SAF-A^ΔRGG^ ^1-7^-GFP for 24 hours. Consistent with our previous work on SAF-A depletion and mutations of the SAP domain, we observed very few changes in gene expression in the first cell cycle after depletion, suggesting that mutation of the SAF-A ATPase and RGG domain mutations had very little impact on overall gene expression (Supplemental Figure 2B-E). These results reinforce the interpretation that gene expression changes that accumulate over time in SAF-A mutant cells are likely to arise as a result of secondary consequences [25].

RPE-1 cells are derived from a somatic cell type and are therefore in the maintenance phase of X-inactivation, during which DNA methylation and histone deacetylation act in parallel with XIST RNA to enforce transcriptional silencing on the Xi [41]. We recently used the haplotype-resolved genome sequence of RPE-1 cells [42] to examine X-linked gene expression using high-throughput sequencing techniques. We found that SAF-A deletion or mutation of the SAP domain did not affect X-linked gene expression, despite having disrupted localization XIST RNA and loss of Polycomb-associated histone modifications [25]. To determine if mutation of the ATPase or RGG domains would dominantly reactivate gene expression from the inactive X chromosome, we performed RNA-seq in cells expressing SAF-A^wt^-GFP, SAF-A^ΔWalker^ ^A^-GFP, SAF-A^ΔWalker^ ^B^-GFP, and SAF-A^ΔRGG^ ^1-7^-GFP, 24 hours after allele replacement (Supplemental Figure 3). To measure allele-specific expression we used the Personal Allele Caller software which was shown to have the highest accuracy and least mapping bias [43]. Examining RNA-seq data from wild-type RPE-1 cells revealed that the vast majority of autosomal genes have equal expression from the maternal and paternal alleles (Supplemental Figure 3A). In contrast, genes located on the X-chromosome exhibit a strong bias with most expression arising from the ‘a’ allele. Consistent with our previous results we can observe duplication of chromosome 10 [42, 44] onto the inactive X chromosome. To determine if mutation of the ATPase or RGG domains resulted in reactivation of genes on the Xi, we used edgeR to compare gene expression from the ‘a’ (active) and ‘b’ (silenced) alleles in RPE-1 cells, and compared to cells expressing SAF-A^ΔWalker^ ^A^-GFP, SAF-A^ΔWalker^ ^B^-GFP, or SAF-A^ΔRGG^ ^1-7^-GFP. Our previous finding was that that genes located on the Xi chromosome exhibited nearly identical gene expression after 24 hours of SAF-A depletion [25]. Similarly, our analysis here shows that none of the SAF-A ATPase or RGG domain mutants caused reactivated gene expression on the Xi (Supplemental Figure 3B-D). We conclude that mutation of the ATPase or RGG domain of SAF-A impacts XIST RNA localization and XIST-dependent organization of PRC1 and PRC2 activity on the Xi, similar to depletion of SAF-A, but does not affect other parallel pathways enforcing transcriptional silencing in the maintenance phase of X inactivation.

### The SAF-A ATPase and RGG domains are required for normal mRNA splicing

Foundational work demonstrated that SAF-A regulates mRNA splicing in multiple organisms and tissue types [8, 17, 45], through a mechanism involving maturation of the U2 snRNP [45]. Additionally, we recently found that acute depletion of SAF-A resulted in a dramatic increase in the missplicing of numerous transcripts. While our findings showed that normal splicing is largely independent of the SAF-A SAP domain [25], it is not currently known whether the ATPase or RGG domains of SAF-A are required for normal mRNA splicing. Therefore, to understand which domains of SAF-A contribute to mRNA splicing, we used rMATS to analyze differential splicing in SAF-A depleted cells, and cells expressing SAF-A^wt^-GFP, SAF-A^ΔWalker^ ^A^-GFP, SAF-A^ΔWalker^ ^B^-GFP, and SAF-A^ΔRGG^ ^1-7^-GFP, using the mRNA-seq data described above. Comparison of the overall number and intersection of misspliced exons revealed that cells expressing SAF-A^ΔWalker^ ^A^-GFP and SAF-A ^ΔRGG^ ^1-7^-GFP showed the greatest number of splicing changes, followed by SAF-A depleted cells, and cells expressing SAF-A^ΔWalker^ ^B^-GFP (Figure 5A and Supplemental Figure 4). Analysis of common misspliced exons revealed the greatest overlap between SAF-A depleted cells and SAF-A^ΔRGG^ ^1-7^-GFP cells, with a lesser overlap observed for SAF-A depleted cells and all other mutants (Figure 5A).

**Figure 5.**
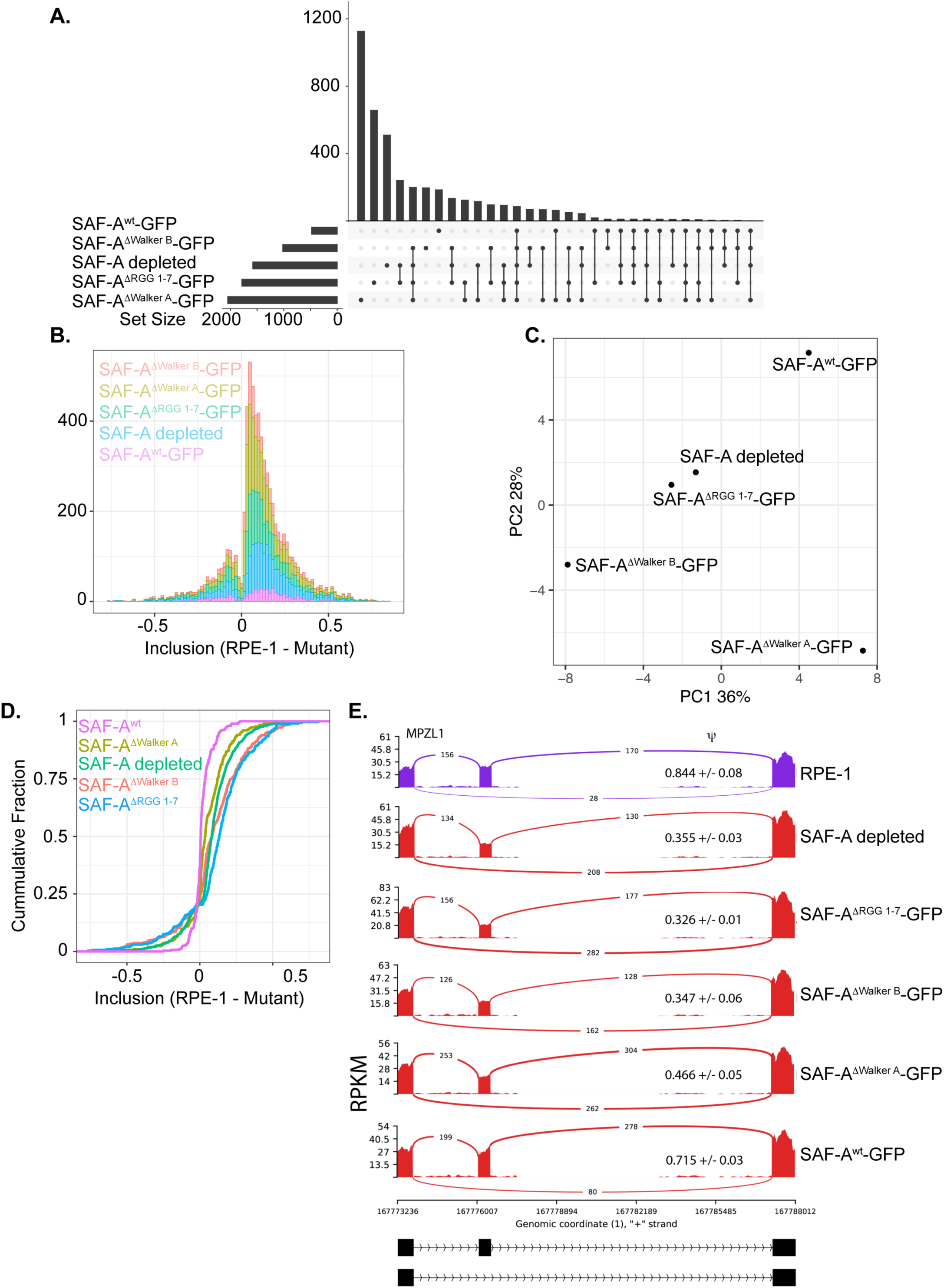
SAF-A depletion leads to widespread changes in mRNA splicing. A. mRNA splicing was evaluated in SAF-A depleted cells (24 hours) using rMATS. Upset plot compares the intersection of all significantly misspliced exons between each condition. B. Histogram plot showing changes in exon inclusion in each SAF-A mutant and the SAF-A depleted cells. C. PCA plot comparing each SAF-A mutant for all common exons assayed. D. CDF plot showing the magnitude of splicing changes for exons that are significantly altered in SAF-A depleted cells for each of the mutants plotted. E. Sashimi plot showing an example altered exon in each mutant.

SAF-A depletion or mutation tended to lead to decreased exon inclusion, although there were examples where loss or mutation of SAF-A led to increased exon inclusion (Figure 5B). Interestingly, the SAF-A^ΔWalker^ ^A^-GFP and SAF-A^ΔWalker^ ^B^-GFP alleles each showed the greatest number of exon inclusion events compared to SAF-A depleted cells, with a similar magnitude of the splicing changes (Figure 5B). SAF-A^ΔRGG^ ^1-7^-GFP cells showed an intermediate level of exon inclusion events, as compared to SAF-A depleted cells and the ATPase domain mutations (Figure 5B). PCA analysis revealed that splicing changes present in SAF-A^ΔRGG^ ^1-7^-GFP cells was most similar to SAF-A depleted cells, while each of the ATPase domain mutants were distinct (Figure 5C).

We then examined the magnitude of exon inclusion for exons that are significantly altered in SAF-A depleted cells and compared these exons in each SAF-A mutant (Figure 5D-E). Interestingly, we identified a clear phenotypic gradient where SAF-A^ΔRGG^ ^1-7^-GFP had significantly stronger alterations in exon inclusion than in SAF-A depleted cells (Figure 5D-E). SAF-A^ΔWalker^ ^B^-GFP was also significantly worse than SAF-A depleted cells, with SAF-A^ΔWalker^ ^A^-GFP having a phenotype that was less severe (Figure 5D-E). To confirm the observed mRNA splicing changes we used qRT-PCR with primer pairs spanning regulated exons. We tested 4 mRNAs exhibiting both increased and decreased exon inclusion in the SAF-A mutants and confirmed the magnitude and direction of all changes predicted by mRNA sequencing (Supplemental Figure 5). Taken together, these results demonstrate that the RGG domain and ability to bind RNA is essential for SAF-A to promote normal mRNA splicing. The ability to hydrolyze ATP is also essential for normal mRNA splicing, while the ability to bind ATP plays a more minor role in promoting splicing. These results suggest that SAF-A SAF-A^ΔRGG^ ^1-7^-GFP and SAF-A^ΔWalker^ ^B^-GFP may be dominant mutants and could sequester SAF-A interacting proteins in nonproductive complexes, leading to strong splicing defects.

## Discussion

In this study we describe the phenotypic consequences of mutation of the SAF-A ATPase and RGG domains in a wide range of essential nuclear RNA biology processes. We used a rapid depletion system in karyotypically stable RPE-1 cells to allow us to observe the immediate consequences of loss or mutation of SAF-A. Importantly, we found that loss of SAF-A RNA-binding, ATP binding or ATP hydrolysis all lead to a loss of cell proliferation, similar to what we observed with mutations to the SAP domain [25]. Interestingly, we observed a phenotypic gradient for each of these mutants when examining RNA splicing. We also observed a differential impact of SAF-A ATP binding and hydrolysis with regard to XIST RNA localization and recruitment of Polycomb-dependent histone marks to the inactive X chromosome. Deletion of the RGG domain resulted in the most severe phenotype in all assays, demonstrating that RNA binding by SAF-A is essential for all reported functions. Mutation of the Walker-B motif also resulted in severe phenotypes in all assays, suggesting that ATP hydrolysis is likely to be important for SAF-A to cycle between different states and or complexes. The SAF-A Walker-B mutant may be trapped in a nonproductive complex and unable to exit this complex. Surprisingly, the Walker-A mutant resulted in the least severe phenotype in most assays. This suggests that the ability to bind ATP is not as important for some SAF-A functions, which may be performed by the DNA and RNA binding activities of SAF-A independent of ATP-induced conformational changes. Collectively, these observations suggest that SAF-A undergoes a complex cycle of RNA binding, DNA binding, ATP binding and ATP hydrolysis and that each of these separate functions are critical for cell proliferation.

Previous work has suggested that SAF-A could function as part of a static ‘nuclear matrix’ [46] or that SAF-A forms oligomeric structures by binding to nascent transcripts and ATP [19]. Our FRAP analysis shows that deletion of the RGG domain leads to a large increase in SAF-A mobility similar to the increase in mobility observed after transcriptional inhibition. This data supports the hypothesis that SAF-A interacts with nascent transcripts throughout the nucleus. However, the rapid dynamics of SAF-A do not support the idea of a static nuclear matrix but rather suggest that SAF-A may be part of a highly dynamic nuclear structure associated with transcriptionally active regions of the genome. Interestingly, our internal deletion of perfect RGG repeats 1-7 dramatically decreases RNA binding both *in vitro* [29] and in cells (Figure 2). This smaller internal RGG deletion exhibits phenotypes different than a deletion of the entire C-terminus [29, 32], suggesting that the extreme C-terminal Q/N-rich domain may have uncharacterized functions that are independent of RNA binding by the RGG domain.

Analysis of SAF-A localization in live cells demonstrated that of all SAF-A domains tested, only the SAP domain is essential for SAF-A recruitment to the Xi territory, while the ATPase and RGG domains are not required. The observation that the RGG domain is dispensable for Xi recruitment was unexpected, since deletion of Xist RNA in murine cells disrupts SAF-A recruitment to the Xi [47], and SAF-A interacts directly with XIST RNA [31]. Because SAF-A^ΔRGG^ ^1-7^-GFP is present on the Xi (Figure 3), we speculate that Xi localization might occur through a protein-protein interaction between SAF-A and part of the XIST ribonucleoprotein (RNP) complex, rather than through XIST RNA binding per se. Once SAF-A is recruited to the XIST RNP, the RNA binding activity contained within SAF-A RGG repeats 1-7 in is required to maintain XIST RNA localization within the Xi chromosome territory.

Our data point to the seventh RGG repeat as being particularly critical for RNA binding activity. RGG repeats 1-6 do not affect RNA binding *in vitro* and are insufficient to bind RNA [29, 30]. However, deletion of RGG repeats 1-7 causes a severe reduction in RNA binding *in vitro and in vivo* [32, this study]. Further, deletion of RGG repeats 1-6 does not affect XIST RNA localization [32], while deletion of RGG repeats 1-7 has a phenotype similar to the null condition (this study).

Collectively, these observations indicate the seventh RGG repeat is critical for both RNA binding and proper XIST RNA localization. Presently, it is not clear how the seventh RGG repeat provides a function distinct from the first six RGG repeats. Further study is needed to understand the unique contribution of the seventh RGG repeat to RNA binding activity.

Analysis of Polycomb-dependent histone modifications on the Xi revealed complex genetic interactions with the SAF-A ATPase and RGG mutations. PRC1 activity directs histone H2AK119ub modification of Xi nucleosomes, and in turn, that modification drives the recruitment of PRC2 to catalyze the histone H3K27me3 modification [40]. Our data suggest SAF-A involvement in directing both PRC1 and PRC2 activity on the Xi. While PRC1 requires the SAF-A SAP domain and RGG domains for optimal activity, PRC2 requires the SAF-A SAP domain, ATP binding, ATP hydrolysis and RGG domains ([32], this study). Based on these observations and the hypothesis that the SAF-A domains are coordinated in a complex reaction cycle, we speculate that SAF-A activity impacts physical interactions between the XIST RNA and protein components of the XIST RNP, thereby influencing chromatin modifications on the Xi.

Our analysis of the ATPase domain demonstrates that both ATP binding and hydrolysis are important for most reported functions of SAF-A. FRAP analysis of Walker-B mutants do not show the existence an immobile complex, arguing against the formation of static SAF-A oligomers in the Walker-B mutant [19]. Interestingly, mutations of either the Walker-A or Walker-B motifs did not alter SAF-A RNA binding in cells, suggesting that RNA binding is primarily controlled by the RGG domain independently of the ATPase domain. Many RNA-binding proteins contain ATPase domains and use conformational changes coupled to ATP binding and hydrolysis to alter the structure of RNAs and RNPs [48, 49]. SAF-A could function as a RNA remodeling enzyme by binding directly to RNA using the ATPase domain, which is supported by the existence of RNA-protein crosslinks in the ATPase domain in several *in vivo* studies [50–52]. *In vitro*, SAF-A has a very slow ATP hydrolysis rate (∼1 ATP hydrolyzed per SAF-A molecule per 5 minutes) that is modestly stimulated by RNA [19]. Most DEAD/DExHD-box helicases have a much higher ATP-hydrolysis rate, suggesting that SAF-A may require a cofactor to stimulate ATP hydrolysis. Additionally, it is not clear whether that SAF-A ATPase domain undergoes conformational changes in response to ATP binding and hydrolysis. Our results clearly show that the SAF-A ATPase domain is required for the normal function of SAF-A in cells, but future work will need to investigate the molecular mechanisms of the SAF-A ATPase domain and how the function of this domain is coupled to the RNA and DNA binding activities of SAF-A.

## Acknowledgements

The authors thank Reito Watanabe for helpful suggestions, and Martha Kaufman-Sharp for assistance with live image acquisition. We thank Daniel Cifuentes for use of the LiCor fluorescent imager. We acknowledge that much of the computational work reported on in this paper was performed on the Shared Computing Cluster which is administered by Boston University’s Research Computing Services. This work was supported by grants from NIGMS to M.B. (5R01GM144352-03, 5R01GM122893-07).

## Figures and Figure Legends

**Supplemental Figure 1.**
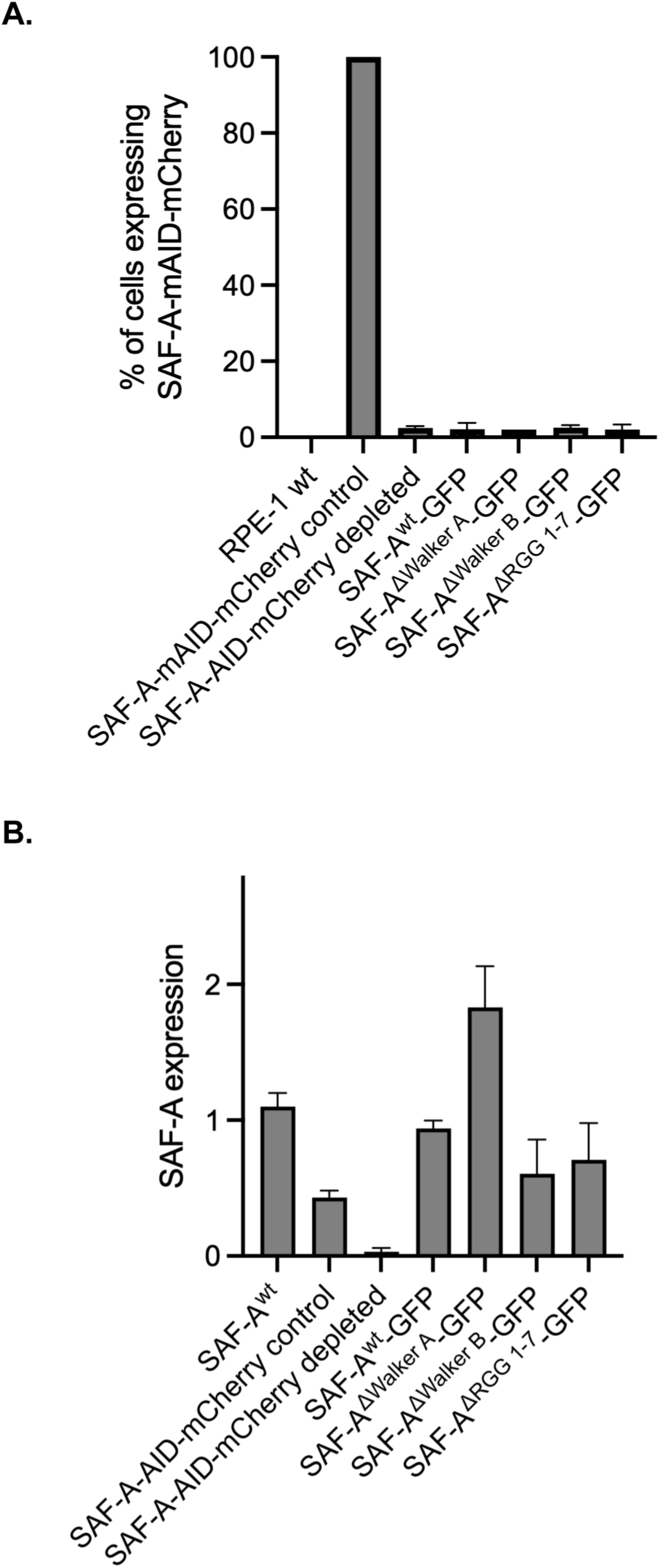
Expression of tagged SAF-A and mutant isoforms. A. Cells were scored for SAF-A-AID-mCherry expression after 24 hours of drug treatment. The graph depicts the average percent of cells (n = 100) expressing the degron allele; error bars depict the SD for ≥2 biological replicates. B. Quantitation of Western blots with the mouse anti-SAF-A 3G6 monoclonal antibody to compare levels of endogenous SAF-A, SAF-A-AID-mCherry -/+ drug treatment, and SAF-A-GFP transgenes. SAF-A expression levels relative to the tubulin loading control in three different extract preparations. Error bars depict SEM of 3 biological replicates. The constitutive expression of SAF-A-AID-mCherry is lower than the endogenous protein, as previously reported [32]. The expression level of all GFP tagged transgenes is within a 2-fold difference compared to the endogenous protein.

**Supplemental Figure 2.**
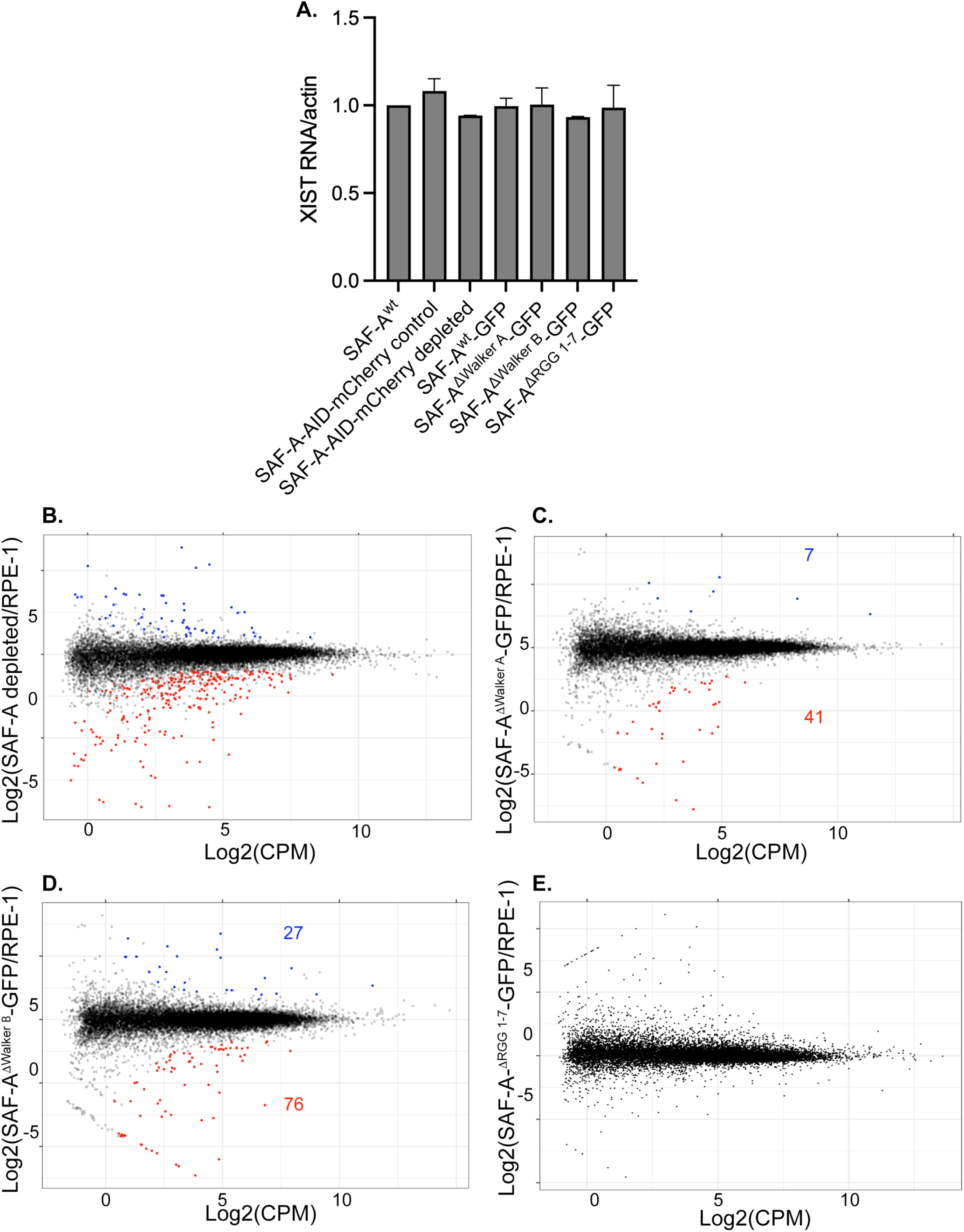
Gene expression in cells expressing SAF-A ATPase and RGG domain mutations. A. qRT-PCR showing XIST RNA gene expression levels relative to actin mRNA. Error bars depict the SD of 2 biological replicates. B-E. Gene expression was evaluated at 24 hours after addition of doxycycline and auxin, using RNA-seq and EdgeR. MD plots depict significantly differentially expressed genes for each mutant (FDR < 0.01). The gene expression profile of cells depleted for SAF-A, or expressing SAF-A alleles SAF-A^ΔWalker-A^-GFP, SAF-A^ΔWalkerB^-GFP, or SAF-A^ΔRGG^ ^1-7^-GFP were compared to RPE-1 cells as indicated on the y axis.

**Supplemental Figure 3.**
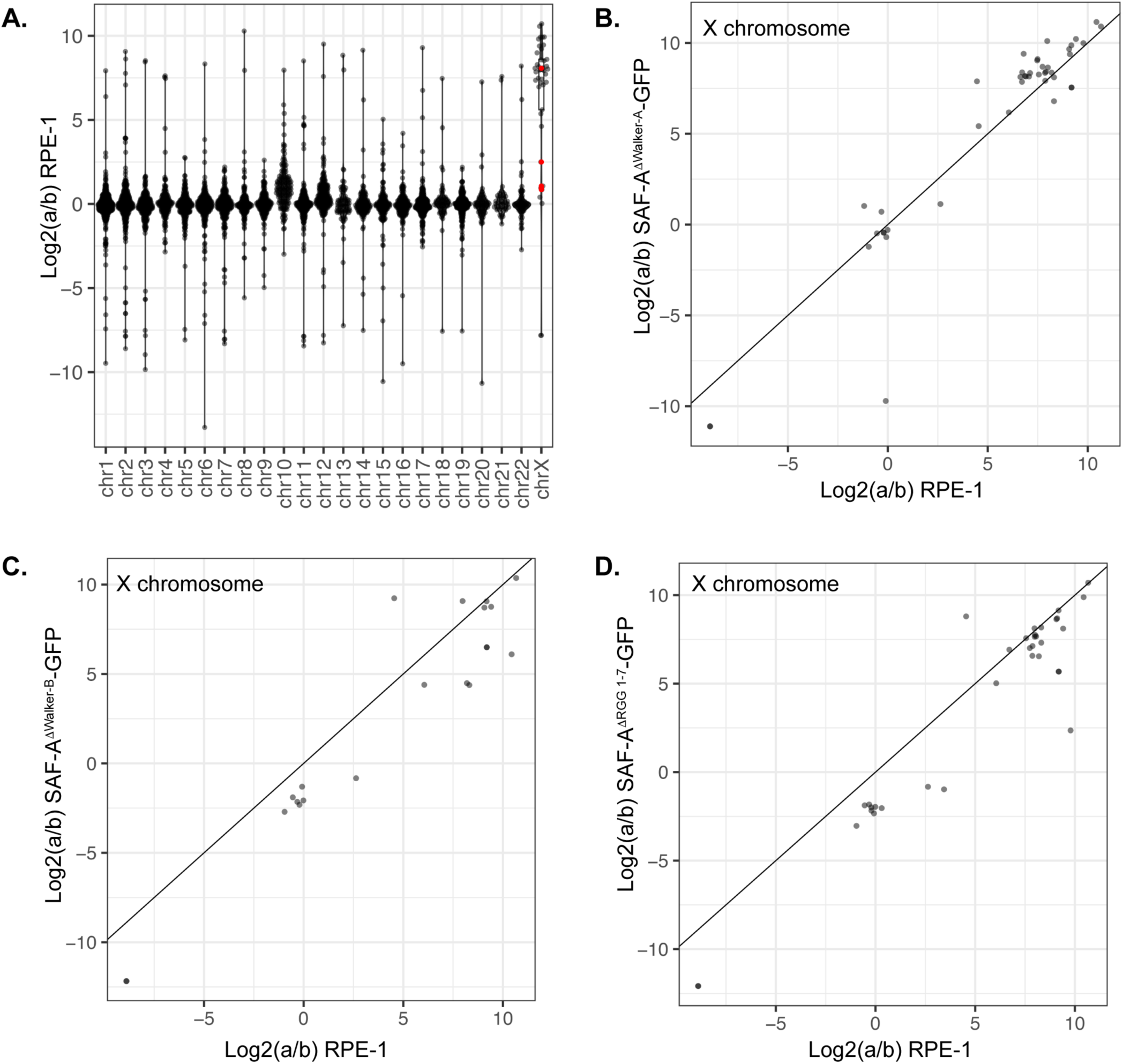
SAF-A depletion does not reactivate gene expression on the inactive X chromosome. A. Allele-specific gene expression was calculated from RNA-seq libraries using a combination of PAC and edgeR in RPE-1 cells. ‘a to ‘b ratios are plotted by gene for each chromosome. B-D. Average a:b ratio for all genes on the X chromosome plotted for RPE-1 and each SAF-A mutant.

**Supplemental Figure 4.**
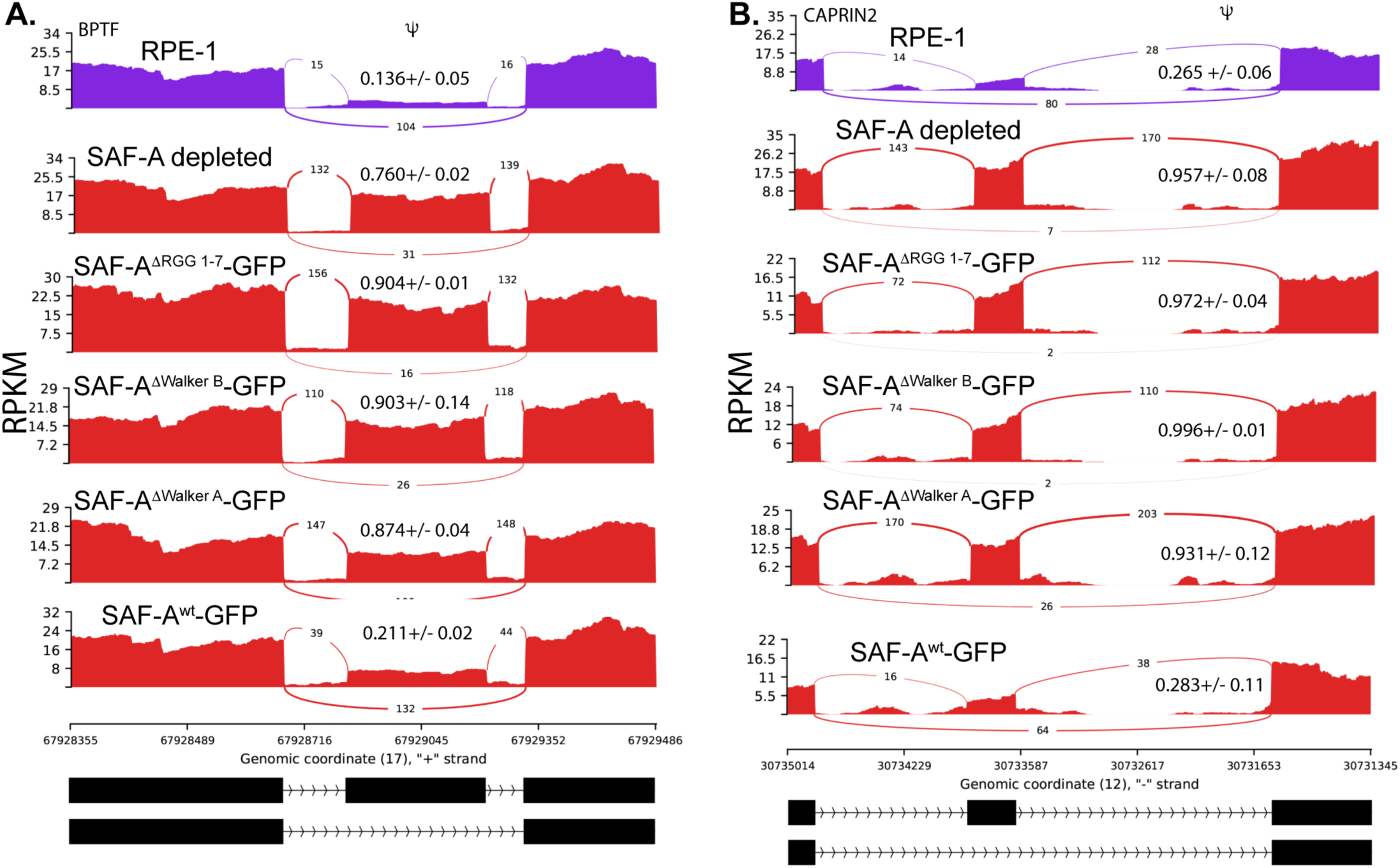
Sashimi plots of additional regulated exons in each SAF-A mutant. A-B. Sashimi plots of SE events in wild-type RPE-1 cells, SAF-A depleted cells, and ATPase and RGG domain mutants.

**Supplemental Figure 5.**
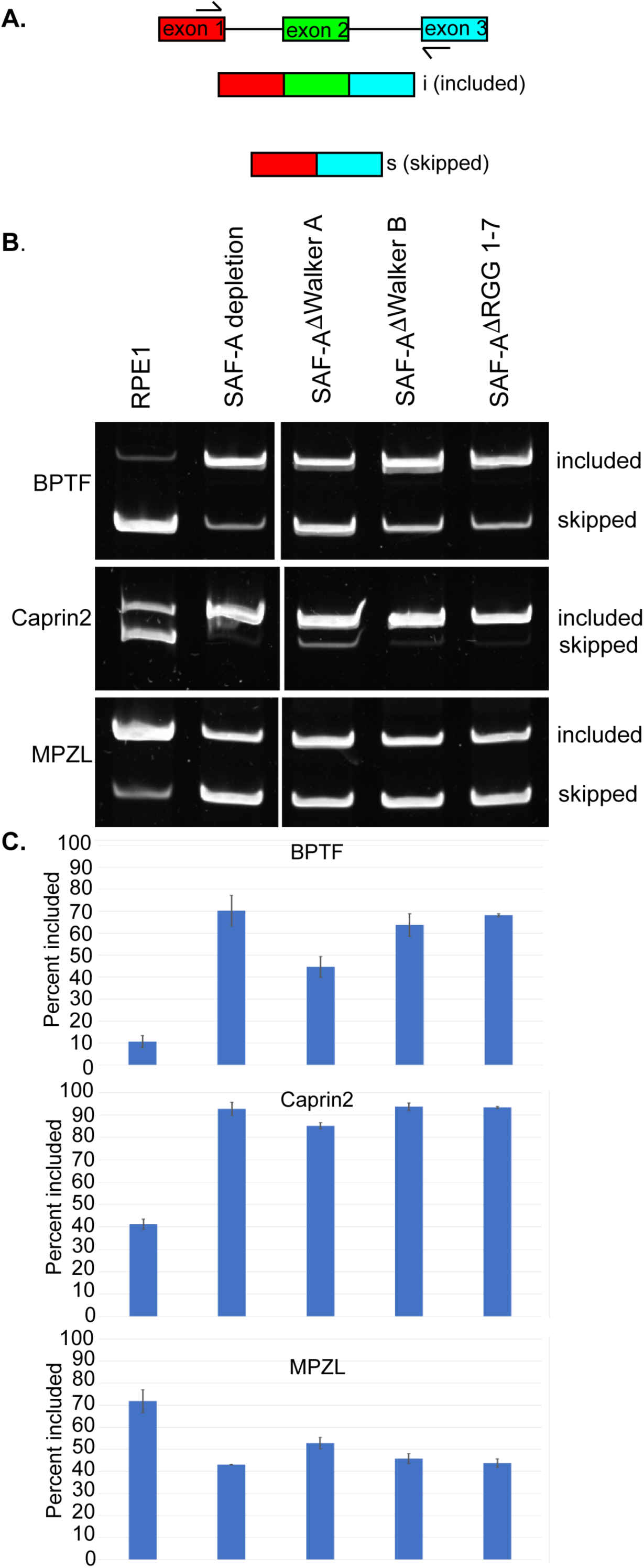
Validation of predicted splicing changes using RT-PCR. A. Illustration depicting included and skipped exons. B. PAGE gel analysis of exon inclusion for 3 different genes with predicted changes in exon inclusion in SAF-A depleted cells. C. Quantitation of percentage spliced in in each mutant from three biological replicates.

## Materials and Methods

### Cell culture and drug treatment

hTERT-immortalized RPE-1 cells were a gift from Brian Chadwick (Florida State University, Tallahassee, FL; ATCC CRL-4000) and were cultured as described [29]. The SAF-A degron cell line in RPE-1 cells has been described [29]. Lentiviruses with SAF-A^wt^-GFP, SAF-A^ΔWalker-A^-GFP, SAF-A^ΔWalker-B^-GFP, and SAF-A^ΔRGG-^GFP were prepared and used to transduce the SAF-A-degron cell line. Individual clones were expanded and validated by immunofluorescent detection of GFP and western blot analysis.

To induce SAF-A depletion +/-SAF-A-GFP expression, cells were treated with 0.5 μg/mL doxycycline (D3447; MilliporeSigma) and 100 μg/mL 3-indole acetic acid (IAA, I-5148 MilliporeSigma) for 24 hours. Control experiments determined that treatment with 0.5 μg/mL doxycycline resulted in SAF-A-GFP expression levels equivalent to the native, untagged protein. For the long-term depletion of SAF-A, media was replaced with fresh drug every 2-3 days.

### Plasmids

GFP-tagged SAF-A alleles were cloned into the lentiviral expression vector pLVX-TetOne-puro (Takara Bio). The lentiviral plasmids pMB1103 (SAF-A^wt^-GFP), pMB1111 (SAF-A^ΔWalker-A^-GFP), pMB1112 (SAF-A^ΔWalker-B^-GFP), pMB1244 (SAF-A^ΔRGG-^GFP), and pMB1316 (SAF-A^ΔSAP^GFP) have been described [29].

### Antibodies

Primary antibodies used in this study were as follows: mouse anti-GFP (sc-9996, Santa Cruz Biotechnology), rabbit anti-mCherry (600-401-p16, Rockland), mouse anti-tubulin (DM1A, MilliporeSigma), GFP booster nanobody-ATTO488 (GBA488, Bulldog Bio), RFP booster nanobody-ATTO 594 (RBA594, Bulldog Bio), mouse anti Ki-67 (610968, BD Biosciences), rabbit anti-H2AK119ub (D27C4, Cell Signaling Technology), mouse anti-H3K27me3 (39537, Active Motif), mouse anti-histone H3 (1B1B2, Cell Signaling Technology), and human anti-PCNA (AK, a gift from Dr. Yoshinari Takasaki (Juntendo University School of Medicine, Japan).

Secondary antibodies used in western blotting were as follows: donkey anti-mouse conjugated to 680RD (925-68072, LiCor) and donkey anti-rabbit conjugated to 680RD (925-68073, LiCor). Secondary antibodies used in immunofluorescent studies were donkey anti-mouse Alexa Fluor 647 (715-605-150, Jackson Immunoresearch), donkey anti-human Alexa Fluor 488 (709-545-149, Jackson Immunoresearch), donkey anti-human Cy3 (709-165-149, Jackson Immunoresearch), donkey anti-rabbit Alexa Fluor 488 (711-545-152, Jackson Immunoresearch), donkey anti-rabbit Cy3 (711-165-152, Jackson Immunoresearch), donkey anti-rabbit Alexa Fluor 647 (Jackson Immunoresearch), and donkey anti-mouse Cy3 (715-165-150, Jackson Immunoresearch).

### Western blot analysis and immunoprecipitation

Cells were grown on 15 cm dishes and washed once with 1X PBS. Cells were incubated for 30 min at 4°C in ice-cold lysis/IP buffer (1 ml per 15-cm plate) containing 25 mM Tris, pH 7.4, 150 mM KCl, 5 mM EDTA, 5 mM MgCl_2_, 1% NP-40, 0.5 mM DTT, and protease inhibitors (Pierce A32955). Cells were collected from the plate and passed several times through a syringe with a 25-gauge needle. Lysates were centrifuged for 30 min at 4°C (∼22,000 ×*g*) to remove insoluble material. The protein concentration of the extract was determined using a Bradford assay. For each cell extract, 10 μg was loaded onto a 4-20% polyacrylamide gel (Bio-Rad 4561096) and run at 15 mA per gel to resolve the different SAF-A protein isoforms. Proteins were transferred to PVDF membrane and probed with antibodies specified above. Blots were exposed using a LI-COR Odyssey 9120 fluorescent imaging system. Blot quantitation was performed using Fiji software.

To immunoprecipitate SAF-A-GFP, cell extracts were prepared as described above, and protein concentration was adjusted to 1 mg/mL. Extracts were incubated with 25 μL pre-equilibrated bead slurry of GFP-Trap Magnetic Agarose (chromotek GTMA020, Bulldog Bio) or Binding Control Magnetic Agarose (chromotek BMAB020) according to the manufacturer’s instructions. After 1 hour incubation at 4°C, immune complexes were collected with a magnet and washed three times with lysis/IP buffer, transferring to a new tube with each wash to reduce background. Total *Drosophila* RNA (a gift of Nelson Lau) was spiked into each immunoprecipitation. RNA was eluted from beads and purified using Trizol. Eluted RNA was used as input to construct RNA-seq libraries as described [25].

### Image acquisition and analysis

All images were acquired using a Nikon A1R confocal microscope equipped with a 60 × 1.4NA Vc lens and laser lines at 405, 488, 562, and 647 nm. Images were collected using Nikon Elements software driving the galvano scanner. Images were acquired as *Z* stacks spaced 0.2 µm apart. Each experiment was repeated in at least two biological replicates. For immunofluorescent detection of SAF-A alleles and Ki-67 (Figure 1E and 1G), we used the same fixation and detection conditions we previously described [29]. For PCNA and immunofluorescent detection of histone H3K27me3 and histone H2AK119ub, we performed permeabilization with CSK buffer (100 mM NaCl, 300 mM sucrose, 3 mM MgCl2, 10 mM PIPES pH 6.8) for 30 seconds, followed by treatment with CSK buffer + 0.5% Triton-X-100, followed by CSK buffer for 30 seconds, and fixation with 4% paraformaldehyde (15710, Electron Microscopy Sciences) buffered with 1X PBS. Our FISH conditions to detect XIST RNA have been published previously [29, 53, 54]. Images were rendered in Fiji software as a projection or single optical slice, as specified in the figure legends.

### Live imaging and FRAP

For live imaging of SAF-A-GFP alleles on the Xi, cells were plated onto fluorodishes (FD35-100, World Precision Instruments) and treated with doxycyline and IAA for 24 hours prior to observation. Hoechst 33342 dye (Molecular Probes) was added just prior to imaging. Cells were imaged on a Nikon A1R confocal microscope equipped with a stage-top incubator and CO_2_ chamber (Tokai Hit). For FRAP analysis, cells were plated as above, except without Hoechst dye. FRAP was performed as follows: the scan area was set to 128 x 128, at a scan zoom size of 8X. 10 images were acquired prior to bleaching. Bleach area was a 3x3 square. Bleaching was accomplished with 80% power of the 488 laser for 1 second. Images were acquired every 65 ms for 30 seconds following bleaching (456 images). For each allele, FRAP movies were collected from at least 25 cells in total on two different days. For experiments involving the transcriptional inhibitor LDC, the drug was added to the culture media 30 minutes prior to imaging. Movies were imported into Fiji and quantitative analysis of FRAP data was performed as described [55]. FRAP curves were plotted in R from average values across all cells analyzed.

### RNA-seq

rRNA depleted samples were converted into libraries using the NEB Next Ultra II RNA library kit. Libraries were sequenced by Novogene using 150 bp paired-end reads.

### RNA-seq analysis

#### Common QC and processing steps

All libraries were sequenced at Novogene using paired-end 150bp reads. Raw fastq files were analyzed for quality, adaptor sequences were trimmed, and duplicate reads were removed using the fastp software suite [56]. Unique, trimmed fastq files were used for all subsequent steps.

#### Differential gene expression analysis

De-duplicated, trimmed reads were aligned to the UCSC hg38 human genome sequence using Rsubread/subread [57]. Reads per gene were quantitated using FeatureCounts in R with Entrez Gene IDs as the reference. Differential gene expression analysis between control and mutant cell lines was performed using edgeR [58]. For the plots in this study we considered genes significant if they exhibited a logFC of >= 1 and FDR value less than 0.01. All MD plots were created using ggplot2 in R.

#### Alternative splicing analysis

De-duplicated, trimmed reads were used as input for rMATS [59] in conjunction with the ENSEMBL hg38 and ENSEMBL transcript annotations. RMATS output files were filtered as follows: first, we only considered splicing events supported by an average of 12.5 junction spanning reads per sample (e.g. for a comparison including two control and two experimental samples we required a total of 50 reads). For determination of the number of significant changes we filtered for a FDR < 0.01. To produce Sashimi plots we used rmats2sashimi.py. To conduct GeneOntology analysis of alternative splicing changes we used the ENSEMBL Gene names of significantly changed exons as input for EnrichR and ClusterProfiler. For analysis of conserved domains altered by alternative splicing we used SpliceTools and batch blast against the NCBI conserved domain database.

#### Allele-specific gene expression analysis

We used the Personalized Allele Caller [43] software in conjunction with the RPE-1 .vcf file [42] of maternal and paternal SNPs to generate counts per gene from each RNA-seq library. To validate the RPE-1 .vcf file with our own batch of RPE-1 cells we sequenced the genome of these cells to a depth of 20X using paired-end 150bp reads. We then ran PAC on this DNA sequencing data using the published .vcf file. We eliminated all SNPs with 0 counts for either allele in the DNA sequencing experiment and all SNPs that exhibited an allele-specific bias of > 1 Standard Deviation from the mean. We used the filtered .vcf file for all subsequent experiments. We filtered the genes to only consider genes with a total of greater than 10 counts per gene. To compare maternal:paternal ratios in different conditions we created average maternal:paternal ratios for each gene from pairs of control and SAF-A depleted/mutant cells. For comparison of RPE-1 to SAF-A depleted cells, this consisted of 6 libraries for each condition. For comparison of RPE-1 to each SAF-A mutant it consisted of 2 libraries for each condition. For the time series of SAF-A depletion, it consisted of 2 libraries for each condition. Averaged ratios were used to create violin plots in R.

#### DGE analysis of gene reactivation on the inactive X chromosome

To determine if genes are reactivated on the inactive X chromosome we used edgeR to calculate a logFC and FDR value for a:b ratios for each gene on the X chromosome. We then compared the logFC (a/b) and FDR values for different genotypes. The number of libraries compared for each condition is described above.

#### ATAC-seq library QC

Sequencing .fastq files were initially QCed using fastp as described above. Reads were aligned to hg38 using bwa-mem. Output .bam files were analyzed for fragment size and genomic location using ATACseqQC [60]. To perform allele-specific ATAC-seq QCed trimmer, deduplicated fastq files were used as input for PAC with the modified .vcf file described above. *Data availability*

All raw reads and summary files are available in GEO under the accession number GSE309811.

### Quantitation of number of XIST particles in SAF-A depleted and SAP mutant cells

3D images of XIST localization in RPE-1, SAF-A depleted cells, and in all SAF-A-GFP cell lines were acquired in 0.2μm z stacks. To quantitate the number of XIST RNA particles in z-stack confocal images, we used a macro implemented in FIJI software. Briefly, we used the DAPI signal to create a mask for nuclei in each image slice after performing a Gausian blur,

Otsu threshold, fill holes and exclude nuclei touching the image edges. Nuclei were then added to the 3D ROI Manager to create linked 3D objects. We then detected XIST RNA particles by segmenting the RNA FISH image using the Otsu method. XIST particles were then added to the 3D ROI manager. We then used the 3D ROI Manager to measure properties of all particles, measure colocalization between all particles, and measure distances between all particles.

Three measurement files were saved for each image. We then used a custom Perl script to summarize the measurements for each nucleus from these three files. Measurements were performed in a batch manner with no user supervision [25].

## Notes

### Competing Interest Statement

The authors have declared no competing interest.

